# HDAC Inhibitors Rescue Multiple Disease-Causing CFTR Variants

**DOI:** 10.1101/399451

**Authors:** Frédéric Anglès, Darren M. Hutt, William E. Balch

**Affiliations:** Scripps Research, Department of Molecular Medicine, 10550 North Torrey Pines Rd, La Jolla, CA, 92037 USA; Skaggs Institute of Chemical Biology, 10550 North Torrey Pines Rd, La Jolla, CA, 92037 USA

## Abstract

Understanding the role of the epigenome in protein misfolding diseases remains a challenge in light of genetic diversity found in the world-wide population revealed by human genome sequencing efforts and the highly variable respond of the disease population to any therapeutic. An ever-growing body of evidence has shown that histone deacetylase (HDAC) inhibitors (HDACi) can have significant benefit in correcting protein misfolding diseases that occur in response to both familial and somatic mutation. Cystic fibrosis (CF) is a familial autosomal recessive disease, caused by genetic diversity in the CF transmembrane conductance regulator (CFTR) gene, a cAMP-dependent chloride channel expressed at the apical plasma membrane of epithelial cells in multiple tissues. The potential utility of HDACi in correcting F508del as well as the over 2000 CF-associated variants remains controversial. To address this concern, we examined the impact of FDA-approved HDACi on the trafficking and function of a panel of CFTR variants. Our data reveal that panobinostat (LBH-589) and romidepsin (FK-228) provide functional correction of class II and III CFTR variants, restoring cell surface chloride channel activity in primary human bronchial epithelial (hBE) cells. We further demonstrate a synergistic effect of these HDACi with Vx809, that together can significantly restore channel activity for multiple CFTR variants. These data suggest that HDACi can serve to level the cellular playing field for correcting CF-causing mutations, a leveling effect that might also extend to other protein misfolding diseases.

## Introduction

CFTR is a multi-membrane spanning polypeptide belonging to the ABC transporter family. It is composed of five functional domains: two nucleotide binding domains (NBD1 & NBD2), two membrane spanning domains (MSD1 & MSD2) and one regulatory domain (RD). CFTR functions as a cAMP sensitive chloride channel at the apical plasma membrane of cells. It is charged with maintaining ion balance and hydration in sweat, intestinal, pancreatic and pulmonary tissues, each providing a unique physiological environment that could affect the synthesis, trafficking and function of this chloride channel [1], differences that are captured through Variation Spatial Profiling (VSP) [2].The biogenesis of CFTR requires trafficking from the endoplasmic reticulum (ER), the first step in the exocytic pathway, through the Golgi to its final destination at the apical cell surface of epithelial cells. The loss of a functional CFTR channel disrupts ion homeostasis resulting in increased mucus viscosity in the airway of the lung [3] and ductal systems of the pancreas and liver and hydration of the intestinal tract [4]. The increased mucus viscosity causes increased risk for inflammation and infection by *P. aeruginosa* in the lung [3] and reduced enzyme secretion in the digestive tract [4].

An analysis of the allele frequency of CF-causing mutations revealed that 70% of patients carry at least one copy of a three base pair deletion leading to the loss of a phenylalanine at position 508 (F508del) in NBD1 [5, 6]. The F508del mutation disrupts the folding of the variant protein, leading to its retention in the ER and clearance by ER-associated degradation (ERAD) [7–13]. While F508del-CFTR is by far the most common CF-associated variant, more than 2.000 disease-causing mutations have been reported in the clinic (www.genet.sickkids.on.ca and www.CFTR2.org), with ∼40% of them predicted to be missense mutations [4]. These mutations are distributed across the entire sequence of the CFTR gene and are grouped into 1 of 6 classes based on their associated functional defect including mutations that lead to a loss of CFTR production (Class I), misfolding and/or premature degradation (Class II), functional impairment (Class III), obstruction of the channel pore (Class IV), a reduction in the amount of CFTR produced (Class V) and destabilization of CFTR at the cell surface (Class VI) [4, 6, 14].

The search for therapeutic solutions that address the genetic diversity responsible for the differential onset and progression of CF disease [2] resulted in the discovery of Lumacaftor (Vx809), a small molecule that correct the trafficking defect associated with the F508del variant and other class II variants [15, 16]. However, it has shown limited and variable clinical value to date [14]. In contrast, a different class of compounds, referred to as ‘potentiators’, such as the compound Ivacaftor, which acts as a small molecule ‘gate opener’, can provide significant improvement in the channel activity of Class III and IV variants (<5% of the CF population) that show variable degrees of trafficking to the cell surface [17, 18]. Ivacaftor is currently approved for clinical use in combination with correctors such as Lumacaftor (Orkambi) and Tezacaftor to partially improve the efficacy of these drugs for Class II variants. They do so, presumably, by improving the channel gating of trafficking corrected pool of class II variants. In some instances, these potentiators can antagonize corrector properties [19]. However, the vast majority of CFTR variants remain largely refractory to current therapeutic approaches. Thus, considerable work is needed to improve patient responses to existing therapeutics, including the heterogeneous clinical response exhibited by patients with the same genotype, effects that might be caused by differences in genetic and epigenetic modifiers.

We now appreciate the importance of epigenetic modifications, such as methylation and acetylation, as key features impacting genetic diversity during development and aging. Epigenomic changes are regulated by Histone Deacetylases (HDACs) and Histone Acetyl Transferases (HATs), as well as by other factors including Histone Methyl Transferases (HMTs), Histone DeMethylases (HDMs), and Bromo-Domain (BD) proteins-together comprising the ‘reader, writer, eraser’ cohort that differentially manage the genome for each individual [20–24]. HATs/HDACs mediate the acetylation balance of histones, which manage the open and closed chromatin states to provide access to transcription factors. They also mediate the acetylation state of numerous non-histone proteins, highlighted by the dynamic acetylome, that includes proteostasis components [1, 25–31] regulating the central heat shock response (HSR) [32] and unfolded protein response (UPR) [33, 34] programs responsible for recovery from protein folding stress [24, 35–41], as observed in CF [42].

How HDAC-regulated epigenomic programs translate genetic diversity in the genome into a functional proteome to contribute to health and disease of each CF individual remains enigmatic. There is now considerable interest in the use of HDAC inhibitors (HDACi) to improve the management of numerous protein folding diseases including CF, Niemann-Pick C1 (NPC1) [43–46], Alpha-1-AntiTrypsin Deficiency (AATD) [47–49], and lysosomal storage disorders [24, 50–65]. Moreover, there is an ever-growing body of evidence that HDACi could play a crucial therapeutic role in somatic or complex diseases including lung fibrosis, Chronic Obstructive Pulmonary Disease (COPD) [66], arthritis, hypertension, septic shock, and neurodegenerative disease including Alzheimer’s, Huntington, and Parkinson’s [67–69]. These results suggest a more general role for HDAC in establishing operational ‘set-points’ in the cell for folding management in response to genetic diversity contributing to human health and disease [2, 70]. This raises the possibility that modulators of HDAC could serve as epigenetic modifiers to manipulate the physiological state of the cell to accommodate the broad range of familial and somatic variation observed in the world-wide CF population.

To address the role of the epigenome in CF biology, we have now examined the impact of a rapidly expanding class of HDAC inhibitors (HDACi) on mitigation of CF etiology [43, 47–49, 71–74]. While the vast majority of these compounds are under investigation as anti-cancer therapeutics where they are used to kill cells, we focus on their potential beneficial effects given the well-recognized role of HDACs in promoting development and differentiation. The human genome encodes 18 HDACs, grouped into 4 classes (I-IV) with a wide range of substrate specificities based on domain structures flanking the highly conserved catalytic site found in each isoform that distinguish the Zn^2+^-dependent class I & II enzymes. HDACi can have both positive and negative effects on gene expression and protein function depending on the Lys/Arg residues affected by the targeted HDAC and the role of these residues in protein function.

It is clear that the epigenetic impact of HDACs represent an important yet unfulfilled opportunity to alter the transcriptional, translational and/or post-translational acetylation balance to improve the function and stability of misfolded proteins driving human disease. Vorinostat, belinostat, panobinostat and romidepsin are four HDACi approved by the US food and drug administration (FDA) to treat cutaneous and peripheral T-cell lymphoma [83–86]. Vorinostat appears to abrogate the phenotypic defects associated with the F508del variant by modulating immune responses [87, 88] and improving the stability and trafficking of F508del-CFTR [89, 90]. While a recent manuscript has reported conflicting data pertaining to the impact of Vorinostat on the F508del variant of CFTR in nasal epithelial cells, the results likely reflect expected cell-specific environments that are differentially sensitive to epigenetic alterations [89]. Given the low and variable response for all CFTR classes to known corrector/potentiator therapeutics, an urgent need exists for general modifiers of drug responses that level the playing field for a given variant within the population.

Herein, we examine the impact of the HDACi belinostat (PXD-101), panobinostat (LBH-589) and romidepsin (FK-228) on the stability, trafficking and function of CFTR variants alone or in combination with the CFTR corrector, Vx809. We provide evidence that these HDACi can correct the trafficking defect associated with class II (ER export defective) & III/IV (gating/channel function defective) CFTR variants and restore cell surface chloride channel activity in primary bronchial epithelial (hBE) cells homozygous for F508del. We observe a synergistic effect of these HDACi when treated in combination with Vx809 for restoring channel activity to therapeutically relevant levels for many CFTR variants.

## Results

### HDAC inhibitor treatment corrects F508del CFTR trafficking

To begin to address the role of the epigenome in managing the cellular proteostasis environment, which determines the fate of disease-associated misfolded proteins, we investigated the impact of three additional FDA approved HDACi, namely PXD-101, LBH-589 and FK-228 (**Figure 1A-C**) [83, 84, 86] on the trafficking and functional correction of F508del CFTR in cell-based CFBE41o- and primary human bronchial epithelial (hBE) cells.

**Figure 1.**
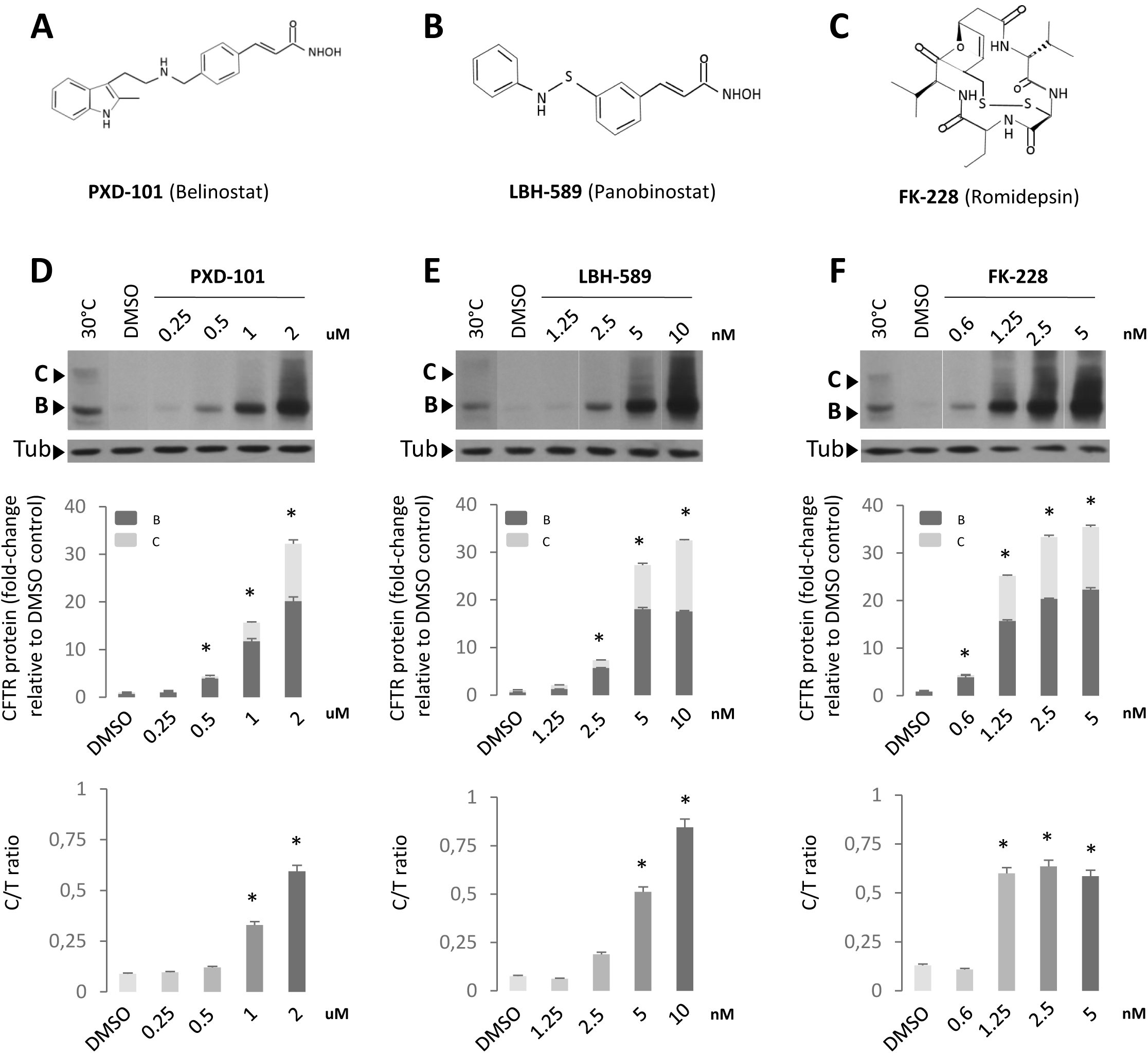
PXD-101, LBH-589 & FK-228 rescue F508del CFTR trafficking in F508del CFBE41o-. Chemical structure of PXD-101 (**A**), LBH-589 (**B**) and FK-228 (**C**). Immunoblot analysis (upper) and quantifications (middle and lower) of CFTR expression following treatment of F508del CFBE41o-cells with different concentration of PXD-101 (**D**), LBH-589 (**E**) and FK-228 (**F**). Data are presented in middle panel as fold change relative to DMSO treatment (mean ± SEM, n ≥ 3). Lower panel represent a quantification of C/T ratio expressed as a % (mean ± SEM, n ≥ 3).

The membrane trafficking of CFTR from the endoplasmic reticulum (ER) to the Golgi compartment can be assessed by monitoring the differential migration of the CFTR glycoforms on SDS-PAGE. The band B fraction represents the N-linked glycosylated ER pool of CFTR (**Figure 1D-F**). The trimming of these glycan chains in the Golgi compartment leads to the slower migration of the post-ER fraction of CFTR on an SDS-PAGE and is represented by the band C fraction (**Figure 1D-F**). To assess the steady state distribution of these glycoforms and their response to HDACi we first utilized CFBE41o-cells, a bronchial epithelial cell line stably expressing the F508del CFTR transgene, referred to herein as CFBE41o-cells. In order to have a reference for the migration of the band B and C glycoforms, we included a loading control of CFBE41o- cells cultured at reduced temperature (30^°^C), which has been shown to cause a partial correction of the trafficking defect associated with F508del CFTR [91]. We observed that treatment with each of the 3 HDACi, PXD-101, LBH-589 and FK-228, caused a dose-dependent improvement in the expression of F508del CFTR (**Figure 1D-F**)- leading to as much as a 30-fold increase in total CFTR relative to that seen with vehicle treatment (**Figure 1D-F**). It is important to note that the increased level of F508del CFTR seen with PXD-101 occurs in the micromolar range, while we obtained similar results with LBH-589 and FK-228 in the nanomolar range.

Given that HDACi’s have been shown to induce cell death [92], we addressed if the observed effects on F508del CFTR were the result of cytotoxic changes in the cellular proteostasis network (PN) leading to the loss of misfolding surveillance in the ER. To this end, we monitored the impact of PXD-101, LBH-589 and FK-228 on the viability of CFBE41o- cells using the LDH cytotoxicity assay, which monitors the integrity of the PM by measuring the amount of the cytosolic enzyme, lactate dehydrogenase (LDH), secreted into the culture media. We did not observe any toxicity with PXD-101 or LBH-589 at doses between 10 pM and 10 μM (**Figure S1**), indicating that these HDACi are not inducing cytotoxicity in the range where we observe changes in CFTR expression. While we did observe toxicity with FK-228 at 10 μM (**Figure S1**), this is 3 orders of magnitude above the doses where we see alterations in CFTR expression. Taken as a whole, these data suggest that the observed effect of HDACi on F508del CFTR is not associated with toxicity associated with treatment of cancer cells.

The increased cellular protein level of F508del CFTR could arise from increased stabilization and/or folding of the polypeptide or could stem from increased transcription. To address this latter possibility, we performed qPCR on HDACi treated CFBE41o- cells at the dose where we observed the maximal increase in CFTR protein levels. We observed a time dependent increase in CFTR mRNA levels starting between 6 and 12 h post HDACi treatment and continuing to increase up to 24 h after treatment (**Figure 2A**).

**Figure 2.**
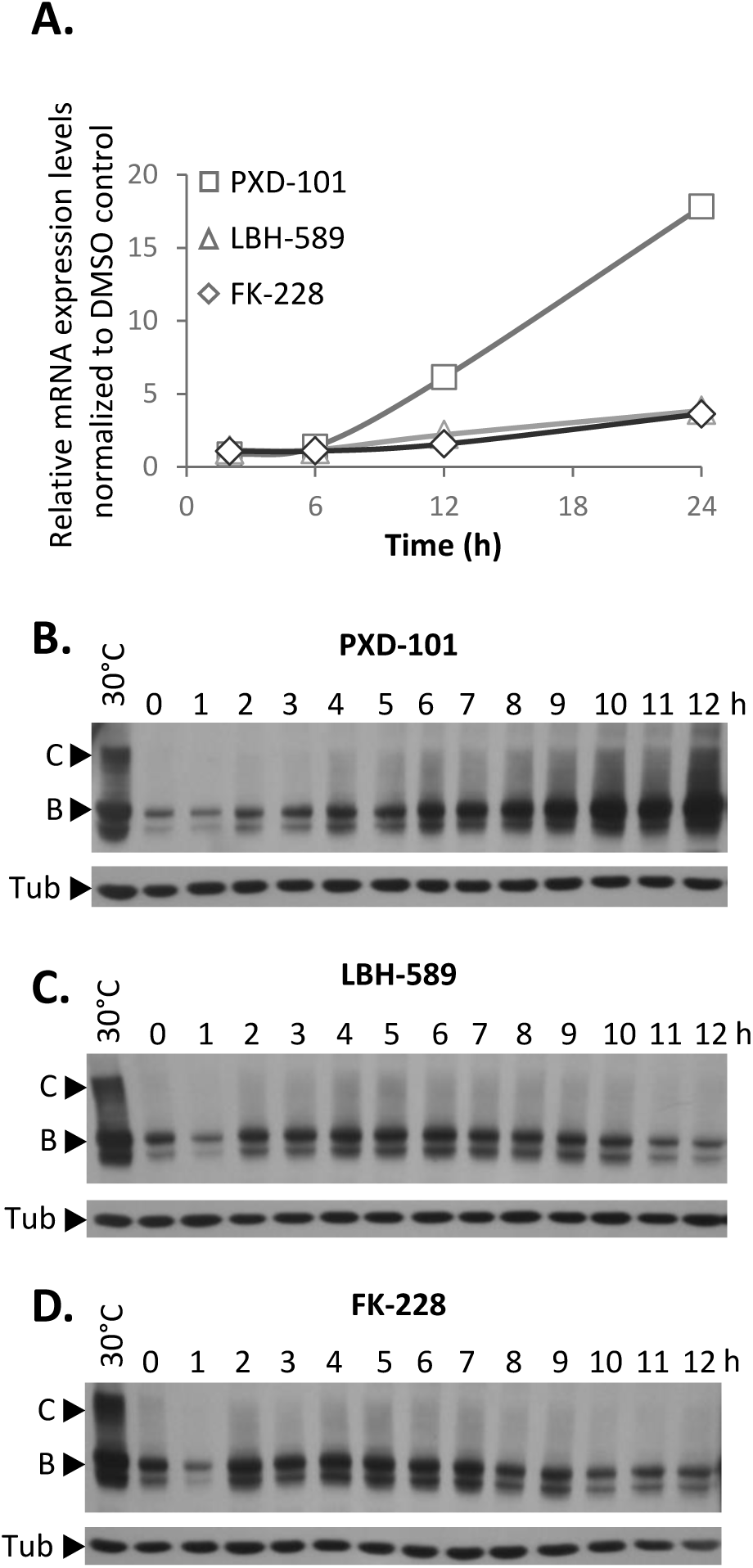
Establishment of F508del CFTR rescue following PXD-101, LBH-589 or FK-228 treatment. (**A**) Quantification of CFTR mRNA level following treatment of F508del CFBE41o- cells with PXD-101, LBH-589 or FK-228 after 2, 6, 12 and 24 hours. mRNA was standardized by quantification of beta-glucuronidase (GUS) mRNA, and all values were expressed relative to GUS (mean ± SEM, n ≥ 3). (**B, C & D**) Immunoblot analysis of CFTR expression following treatment of F508del CFBE41o- cells with 2 µM PXD-101, 10 nM LBH- 589 and 5 nM FK-228 respectively.

To address if the protein expression levels tracked with changes in mRNA levels we next monitored the changes in F508del CFTR protein levels over the first 12 h of exposure to PXD-101, LBH-589 and FK-228. For PXD-101. We observed no changes in CFTR mRNA level in the first 6h of exposure (**Figure 2A**), however we do detect changes in the protein levels of F508del CFTR starting as early as 2 h and continuing to increase up to 24 h (**Figure 2B & Figure S2A**). Treatment of CFBE41o- cells with LBH-589 and FK-228 also did not induce statistically significant changes in the mRNA levels of F508del CFTR in the first 12 h (**Figure 2A**) but did cause an increase in the protein levels of F508del CFTR (**Figure 2C-D & Figure S2B-C**). Interestingly, the altered protein levels seen in response to LBH-589 and FK- 228 appear to be biphasic, as we observed a decrease in F508del CFTR protein levels occurring between 6 and 12 h post-treatment (**Figure 2C & D**) and returning to elevated levels at 24 h (**Figure 1E & F**). These data show that early in the HDACi dosing period, transcriptional and/or translational changes impact the stability of the F508del variant prior to changes in CFTR transcription itself. These data suggest that the observed changes are not solely the result of increased CFTR expression. Additionally, we observed that all 3 HDACi induced a similar 30-fold increase in F508del CFTR protein levels relative to that seen with DMSO treatment in CFBE41o- cells (**Figure 1D-F**), yet the mRNA changes induced by PXD- 101 (18-fold) are significantly greater than that seen with either LBH-589 or FK-228 (3-fold), suggesting that other cellular factors are responding to these HDACi to impact the stability and/or folding of the resulting F508del CFTR protein.

Increased levels of CFTR are not necessarily indicative of improved trafficking of CF-causing disease variant [93]. We therefore performed a more detailed analysis of our data, which revealed that all 3 HDACi cause a dose-dependent improvement in the level of the band C glycoform (**Figure 1D-F**), suggesting that they are providing correction of the trafficking defect associated with F508del CFTR. In order to assess if the HDACi treatment of CFBE41o- cells is increasing the trafficking efficiency of F508del CFTR, we also calculated the ratio of band C obtained relative to the amount of total CFTR (referred to as the C/T ratio), a trafficking index which determines whether the increase in band C is proportional to the increase in the ER pool generated or whether a given treatment has altered the fractional distribution of the CFTR glycoforms.

An analysis of the C/T ratios for the early response of the F508del CFTR protein to PXD-101, LBH-589 and FK-228 revealed differing patterns of response to each HDACi. We did observe an increase in the C/T ratio with PXD-101 but only following changes in CFTR mRNA levels (**Figure 2A & Figure S2A**). The response to LBH-589, revealed no changes in the C/T ratio in the first 12h of the dosing regimen (**Figure 2C & Figure S2B**) suggesting that the improved trafficking occurred much later in the dosing period (**Figure 1E**). Conversely, the treatment with FK-228 induced a biphasic improvement in F508del CFTR trafficking, with a significant increase in C/T observed between the 2 and 7 h mark (**Figure 2D & Figure S2C**) and again at the 24 h mark (**Figure 1F**). Taken as a whole, these data suggest that the HDACi are in fact not only increasing the level of total F508del CFTR, but also improving the trafficking efficiency of this CF-causing variant.

### HDAC inhibitor restore a functional F508del CFTR at cell surface

We next determined if the increased band C fraction represented a functional cell surface-localized chloride channel. In order to assess the functional status of F508del CFTR in CFBE41o- cells treated with PXD-101, LBH-589 or FK-228, we used CFBE41o- cells stably expressing the halide-sensitive YFP-H148Q/I152L, a YFP variant whose fluorescence can be quenched in response to iodide influx entering the cell through a functional, cell surface localized CFTR channel [94]. Treatment with the PXD-101, LBH-589 or FK-228 caused a dose-dependent increase in YFP quenching in CFBE41o- cells (**Figure 3A-C**) at the same doses where we observed correction of the trafficking defect of F508del (**Figure 1D-F**). CFBE41o- cells treated with 2 μM PXD-101, 10nM LBH-589 and 1.25 nM FK-228 produced similar level of YFP quenching to that seen in WT CFTR expressing CFBE41o- cells **Figure 3D** (WT activity)). This result indicates that the HDACi-corrected F508del CFTR is functional and the amount of F508del CFTR found at the cell surface is sufficient to restore wild-type like activity in CFBE41o- cells.

**Figure 3.**
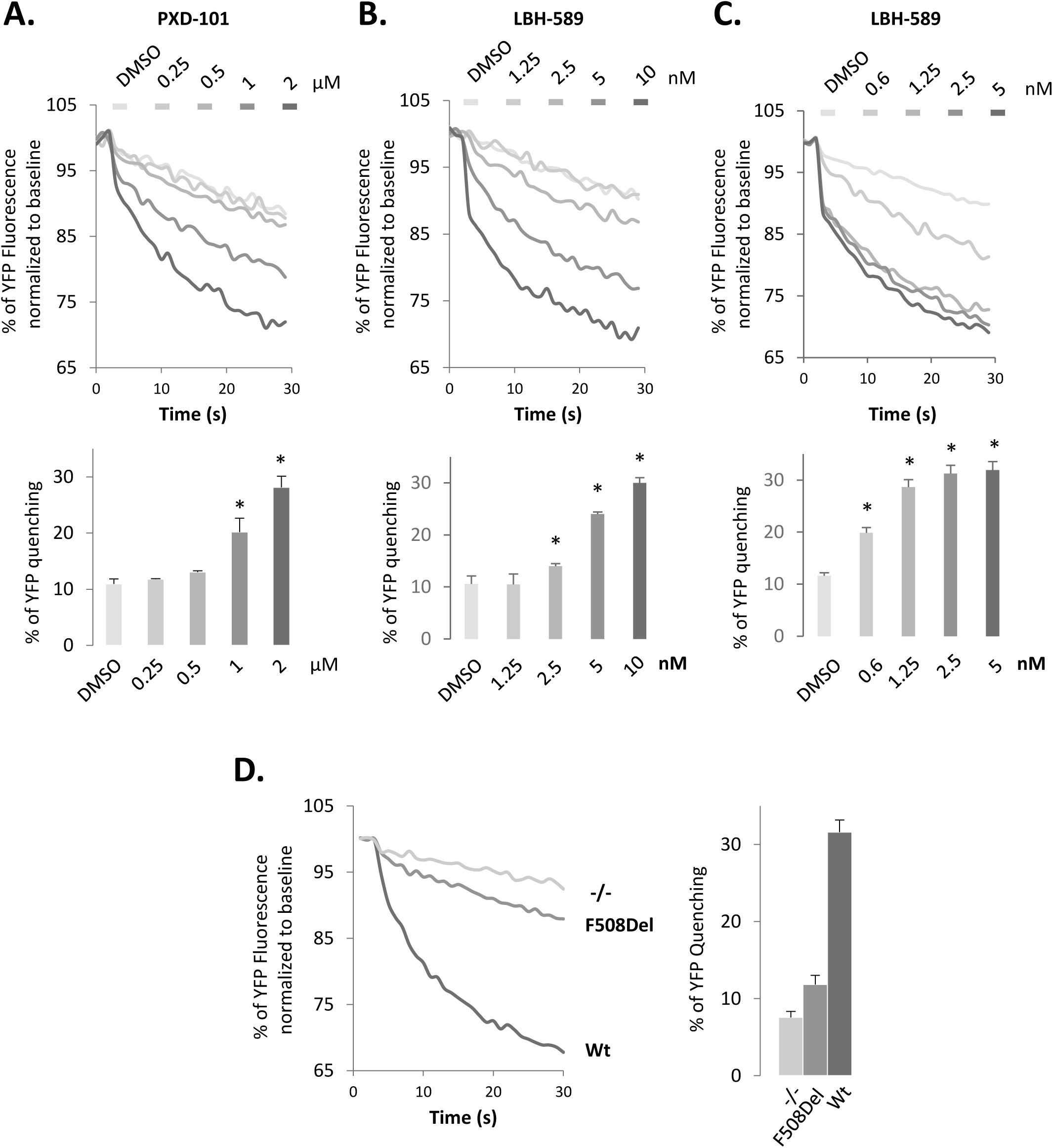
PXD-101, LBH-589 & FK-228 rescue F508del CFTR function in F508del CFBE41o-. Representative FLIPR traces (upper) and quantification (lower) of YFP- quenching following treatment of F508del CFBE41o-YFP cells with different concentrations of PXD-101 (**A**), LBH-589 (**B**) and FK-228 (**C**). Data are presented as % relative to baseline (mean ± SEM, n ≥ 3). In all panels, * indicates significant differences (p < 0.05) relative to DMSO treatment as determined by two-tailed t-test. (**D**) Representative FLIPR traces (left) and quantification (right) of YFP-quenching of CFBE41o-YFP, F508del CFBE41o-YFP and WT-HBE41o-YFP cells. Data are presented as % relative to baseline (mean ± SEM, n ≥ 3).

### HDAC inhibitors synergize with Vx809 to correct F508del CFTR

While numerous efforts have been made to identify small molecule correctors of the F508del variant of CFTR, the best characterized is Lumacaftor (Vx809) [15, 95]. Treatment of CFBE41o- cells with Vx809 resulted in a 2-fold increase in the C/T ratio of F508del CFTR without causing a global increase in total F508del CFTR protein level (**Figure 4A**). As expected, the Vx809-mediated correction of the trafficking defect of F508del CFTR correlated with a very modest 50% increase over the basal channel activity seen in F508del-expressing cells (**Figure 4B-D**).

**Figure 4.**
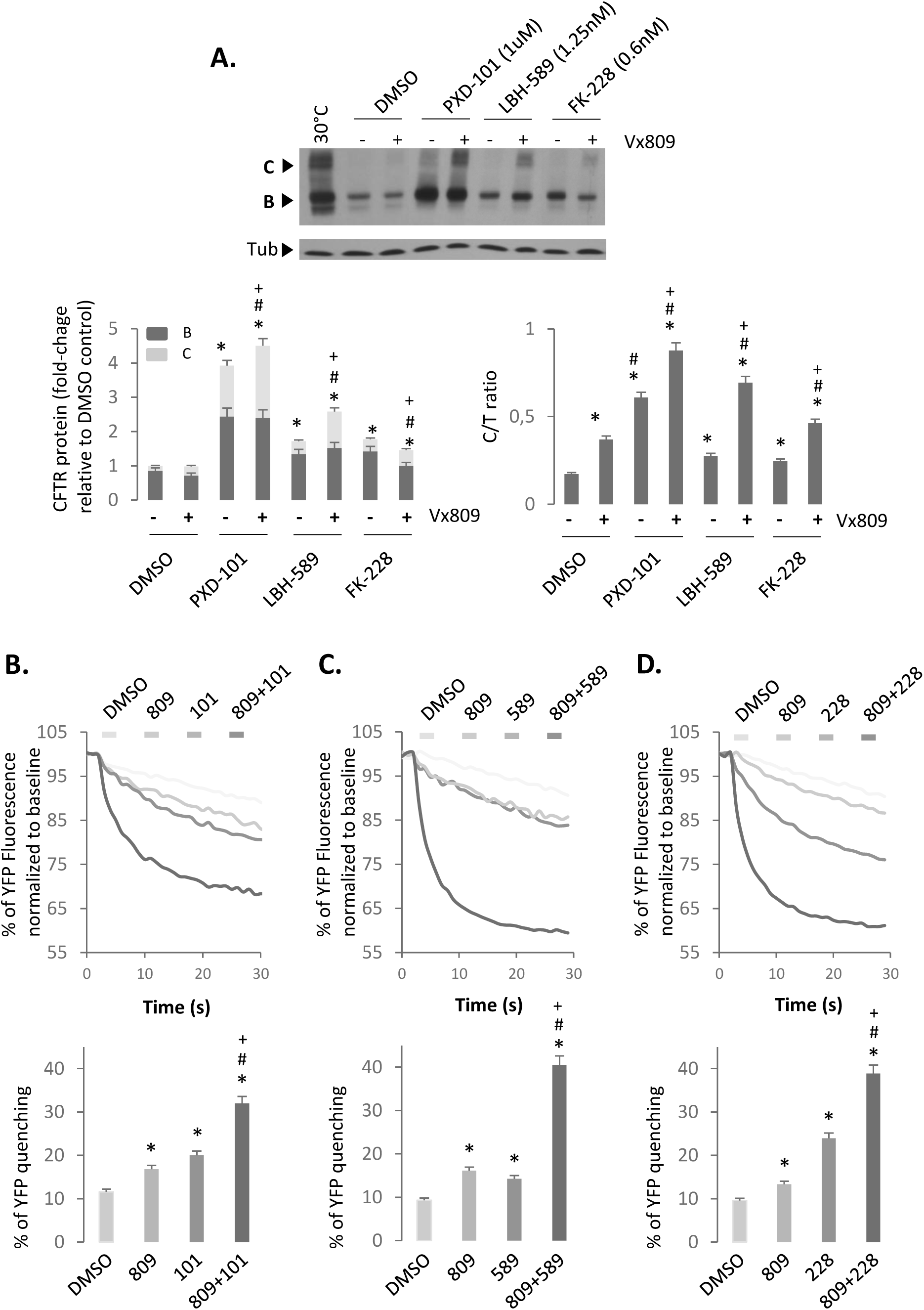
Synergistic effect of PXD-101, LBH-589 or FK-228 in combination with Vx809. (**A**) Immunoblot analysis (upper) and quantification (lower) of CFTR expression following treatment of F508del CFBE41o- cells with 1 µM of PXD-101, 1.25 nM LBH-589 or 0.6 nM FK-228 with or without 3 µM of Vx809. Data are presented as fold change relative to DMSO treatment (lower left) or as a % of C/T ratio (mean ± SEM, n ≥ 3) (lower right). Representative FLIPR traces (upper) and quantification (lower) of YFP-quenching following treatment of F508del CFBE41o-YFP cells with 1 µM of PXD-101 (**B**), 1.25 nM LBH-589 (**C**) or 0.6 nM FK- 228 (**D**) with or without 3 µM of Vx809. Data are presented as % relative to baseline (mean ± SEM, n ≥ 3). Abbreviations: 809 (Vx809), 101 (PXD-101), 589 (LBH-589) and 228 (FK-228). In all panels, * and # indicate significant differences (p < 0.05) relative to DMSO and Vx809 treatment respectively as determined by two-tailed t-test. + indicates significant difference between the same HDACi treatment with or without Vx809.

The inability of Vx809 to provide statistically significant improvement in patients homozygous for the F508del variant initially spelled failure for this drug [96], however recent efforts have shown clinical benefit of Vx809 when combined with the FDA-approved small molecule potentiator, Ivacaftor (Vx770) in patients carrying at least one F508del allele [97–99] in the form of Orkami and more recently as the improved Lumacaftor scaffold Tezacaftor [100]. These data highlight that Vx809, while able to weakly correct the trafficking defect associated with F508del, requires additional “help” to overcome the threshold level of PM-resident F508del to achieve global improvement in cellular chloride channel activity or to potentiate the corrected channel localized in the PM.

In light of these observations, we investigated the effect of combining Vx809 with PXD-101, LBH-589 or FK-228 on F508del CFTR. Considering the potency of the HDACi at maximal dosing (**Figure 1D-F**), we utilized doses of PXD-101 (1 μM), LBH-589 (2.5 nM) and FK-228 (0.6 nM) where the corrective benefit was just beginning to be observed to allow for potential additive or synergistic effects to be detected. When we combined these lower doses of HDACi with a treatment of 3 μM Vx809 we observed an additive effect on the level of band C glycoform observed relative to the effect seen with Vx809, PXD-101, LBH-589 and FK-228 alone (**Figure 4A**). The combined treatment of PXD-101 or FK-228 and Vx809 resulted in an additive effect on the trafficking efficiency of F508del CFTR (**Figure 4A**). However, combining Vx809 with LBH-589 resulted in a synergistic improvement in the trafficking efficiency of the F508del variant (**Figure 4A**).

Given the impact of these combinatorial treatments on improving the level of band C correction for the F508del variant, we next addressed the functionality of the accumulated band C using the YFP-quenching assay. We observed a synergistic improvement in the amount of functional F508del CFTR in cells treated with Vx809 in combination with PXD-101, LBH-589 or FK-228 compared to that seen with any of the treatments alone (**Figure 4B-D**). Interestingly, we observed a greater increase in F508del activity in cells treated with Vx809 combined with either LBH-589 or FK-228 than with PXD-101 (**Figure 4B-D**) despite detecting a more pronounced increase in total CFTR and band C with the PXD-101/Vx809 combination treatment than with the former 2 combinations (**Figure 4A**). Additionally, we observed additivity in the trafficking by combining Vx809 with either PXD-101 or FK-228, yet observed synergistic increases in activity of the corrected F508del fraction, suggesting that the combinatorial treatments are altering the channel properties of the F508del variant such that the measurable activity is no longer a direct correlation of its band C expression. These results suggest that the combined treatment with LBH-589 and FK-228 provide a more effective improvement in F508del CFTR stability, trafficking, folding and/or function than PXD-101 alone.

### HDACi rescue F508del CFTR in primary airway cells

While CFBE410- cells are a commonly used cellular model to study the folding, trafficking and function of CFTR variants and have proven useful in the identification of small molecule therapeutics and targets to treat CFTR phenotypic defects, they express CFTR from a transgene driven by a viral promoter, which fails to recapitulate CFTR biology and its endogenous promoter activity that are likely to have tissue specific responses triggering variable levels of expression and/or the maladaptive stress response (MSR) that is known to contribute to the etiology of disease and its corrective potential through proteostasis modulation [101]. Consistent with this view, a recent report indicated that Vorinostat cannot correct nasal epithelial cell conductance when placed in transwell culture, conditions which recapitulate the nasal respiratory environment [89].

In order to address the potential therapeutic benefit of PXD-101, LBH-589 and FK- 228 in correcting the defects associated with the F508del variant in the lung, we tested their ability to provide functional correction of endogenously expressed F508del CFTR in primary human bronchial airway cells (hBE) isolated from the lung of a CF patient homozygous for the F508del mutation. The treatment of hBE cells with Vx809 results in a modest 25% increase in the band B glycoform of F508del CFTR as well as a 300% increase in the post-ER band C fraction, confirming the ability of Vx809 to modestly correct the trafficking of F508del CFTR (**Figure 5A**). The Vx809-corrected band C fraction represents a functional F508del CFTR chloride channel as we observe a 3.5-fold increase in the forskolin (Fsk) and Genistein (Gen) stimulated short-circuit current (I_sc_) measurements in hBE cells (**Figure 5B-D & F)**. Following treatment with Vx809, the channel activity reaches a level of 16.2 μAmp/cm^2^ a level representing ∼25% of wild type channel activity.

**Figure 5.**
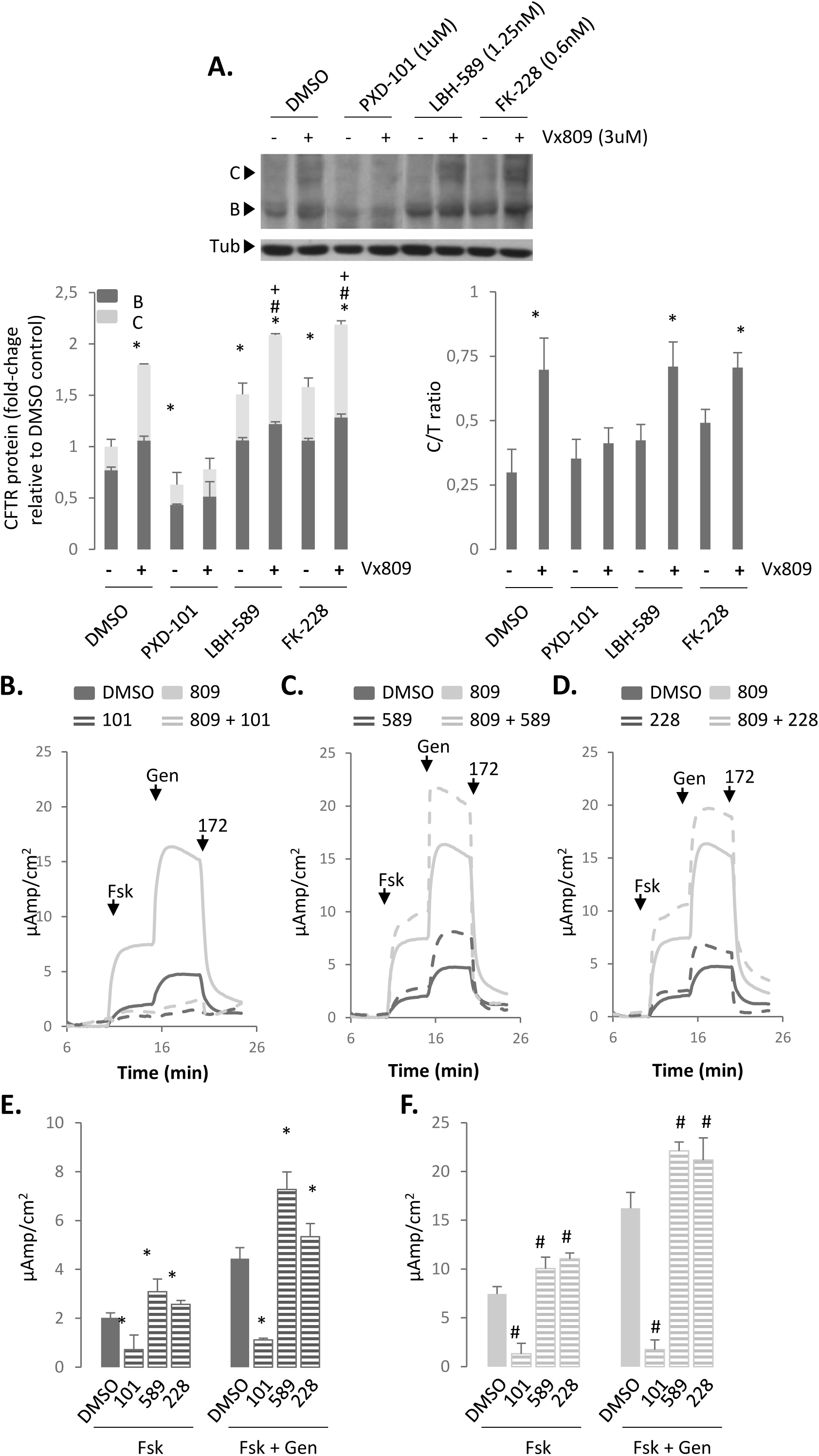
Impact of PXD-101, LBH-589 or FK-228 alone or in combination with Vx809 on F508del CFTR in hBE airway cells homozygous for F508del. (**A**) Immunoblot analysis (upper) and quantification (lower) of CFTR expression following treatment of F508del/F508del hBE primary cells with 2 µM of PXD-101, 5 nM LBH-589 or 5 nM FK-228 with or without 3 µM of Vx809. Data are presented as fold change relative to DMSO treatment (lower left) or as a % of C/T ratio (mean ± SEM, n ≥ 3) (lower right). Representative short-circuit current (Isc) traces for F508del/F508del hBE following treatment with 2 µM PXD-101 (**B**), 5 nM LBH-589 (**C**) and 5 nM FK-228 (**D**), with or without 3 µM of Vx809. Quantification of Isc for F508del/F508del hBE following treatment with 2 µM PXD-101, 5 nM LBH-589 and 5nM FK-228 without (**E**) or with (**F**) 3 µM of Vx809 respectively (mean ± SEM, n ≥ 3). Abbreviations: 809 (Vx809), 101 (PXD-101), 589 (LBH-589) and 228 (FK-228). In all panels, * and # indicate significant differences (p < 0.05) relative to DMSO and Vx809 treatment, respectively, as determined by two-tailed t-test.

In stark opposition to the observations made in CFBE41o- cells, PXD-101 treatment of hBE cells resulted in a destabilization of the F508del CFTR protein, which could not be overcome by combining the treatment with Vx809 **(Figure 5A**). A measurement of I_sc_ for hBE cells treated with PXD-101, either alone or in combination with Vx809 revealed a complete loss of Fsk/Gen-stimulated chloride channel activity (**Figure 5B, E, F**). Conversely, LBH-589 and FK-228 both induced a stabilization of the band B pool and improved trafficking to band C (**Figure 5A**). The increase in the post-ER pool of F508del CFTR induced by LBH-589 and FK-228 represented functional chloride channels as determined by I_sc_ measurements, where we observed an ∼50% and 25% increase in the basal channel activity, respectively. These effects significantly impact the effect of Vx809. HDACi-mediated stabilization of CFTR further improved the Vx809 sensitive trafficking of F508del CFTR leading to a statistically significant improvement in the C/T trafficking index (**Figure 5A**). Moreover, combining HDACi and Vx809 in the hBE cell model improved the cell surface channel activity seen with Vx809 alone leading to correction of the F508del CFTR functional defect to a level representing 37% and 33% of WT CFTR channel activity (**Figure 5C-F**), a level considered to provide significant benefit in the clinic [102, 103]. These data support the potential therapeutic benefit of LBH-589 and FK-228 both as solo treatments or in combination with small molecule correctors such as Vx809.

### LBH-589 and FK-228 modulate CFTR2 variant stability, trafficking and activity

While F508del CFTR is the most common CF-causing mutation, there are currently more than 2000 disease-associated variants reported in the clinic (www.genet.sickkids.on.ca and www.CFTR2.org) [4]. A large-scale study, which examined the most common mutations (allele frequency ≥ 0.01%) representing ∼96% of the CF population, identified 67 missense CF-causing mutations, with nearly half of these variants exhibiting a trafficking defect (class II). We characterized the impact of HDACi on 9 class II variants: P67L, G85E, E92K, S492F, F508del, R560T, L1077P, M1101K and N1303K, as well as the class III G551D variant, the 3^rd^ most common CF-causing mutation in the patient population [6], which exhibits WT-like trafficking but is characterized by a gating defect. The class II mutations are distributed throughout the CFTR polypeptide (**Figure 6A**), and predicted to induce different structural changes and their respective phenotypic responses [2].

**Figure 6.**
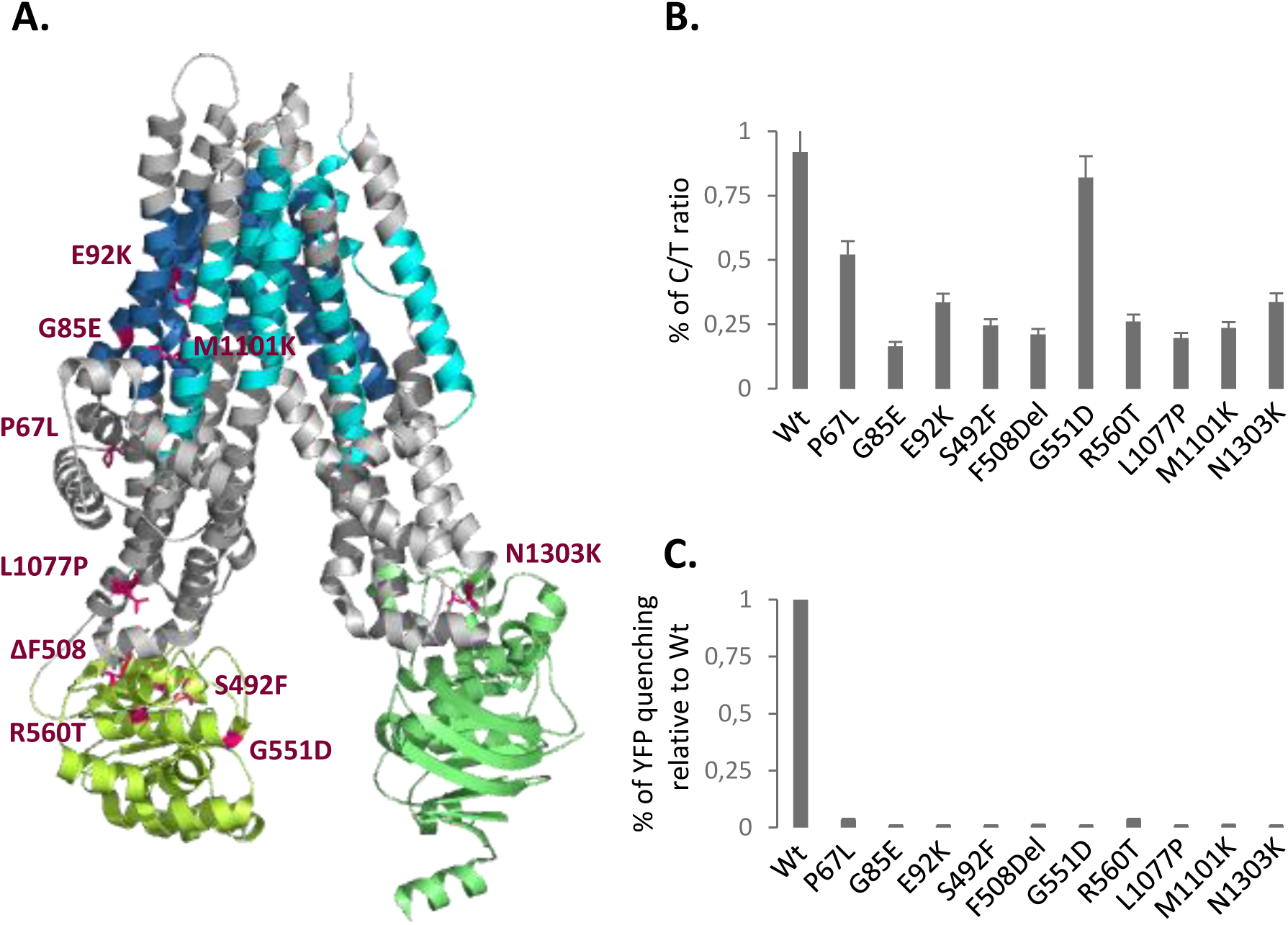
Characterization of CFTR2 variant trafficking. (**A**) Cryo-EM model of human CFTR [114] (blue: TMs of MSD1, yellow-green: NBD1, cyan: TMs of MSD2 and green: NBD2) showing the positions of the selected CFTR2 mutations in Magenta. (**B**) Quantification of C/T ratios following adenovirus transduction of CFTR2 variants into CFBE41o- cells. C/T ratios are expressed as a % (mean ± SEM, n ≥ 3). (**C**) Quantification of YFP-quenching following adenovirus transduction of CFTR2 variants into CFBE41o-YFP cells. Data are presented as % normalized to WT (mean ± SEM, n ≥ 3).

To perform a side-by-side comparison of the different CF variants, we utilized parental CFBE41o- cells, which are genotypically F508del/F508del, and do not express detectable CFTR mRNA or protein. The choice of these CFBE41o- null cell lines provide a cellular environment optimized for the expression, folding, trafficking and functional regulation of CFTR [104]. To characterize the 10 variants described above in response to HDACi in the presence or absence of Lumacaftor, we transduced the CFBE41o- null cells with adenoviral particles carrying the different CFTR variant cDNAs. While the transient expression level of the F508del variant in CFBE41o- null cells was much lower than seen in CFBE41o- stable cell lines, the stability and trafficking of the variants observed is consistent with what has been previously reported for these variants [105, 106] (**Figure 6B & 7**).

Using 5 nM LBH-589 and 2.5 nM FK-228, doses shown to provide maximal correction of F508del CFTR in CFBE41o- cells, we observed a significant improvement in the stabilization of band B for all CFTR2 variants tested as well as correction to band C (**Figure 7A-K**). We also observed improvement in the trafficking efficiency for all variants tested (**Figure 7A-K**). We observed that P67L, E92K, F508del and M1101K-CFTR modestly responded to the Vx809 treatment **(Figure 7A-J**). The combination treatment of Vx809 with LBH-589 or FK-228, where the HDACi-mediated increase in CFTR expression level for the P67L and E92K variants resulted in a much larger pool of CFTR that can be corrected by Vx809 (**Figure 7A & C**), with a significant increase in the C/T ratio over that seen with LBH-589 or FK-228 alone (**Figure 7K**). Not all variants responded equally as other tested variants failed to show an additive or synergistic effect in response to the combined treatment of HDACi and Vx809 possibly reflecting the impact of LBH-589 and FK-228, whose effects alone would mask the weak impact of Vx809 observed when treated alone. Interestingly, we observed that both LBH-589 and FK-228 increased the amount of G551D band C without affecting the C/T trafficking index, consistent with the observation that CFTR variants that do not exhibit trafficking defects already exhibit efficient ER export.

**Figure 7.**
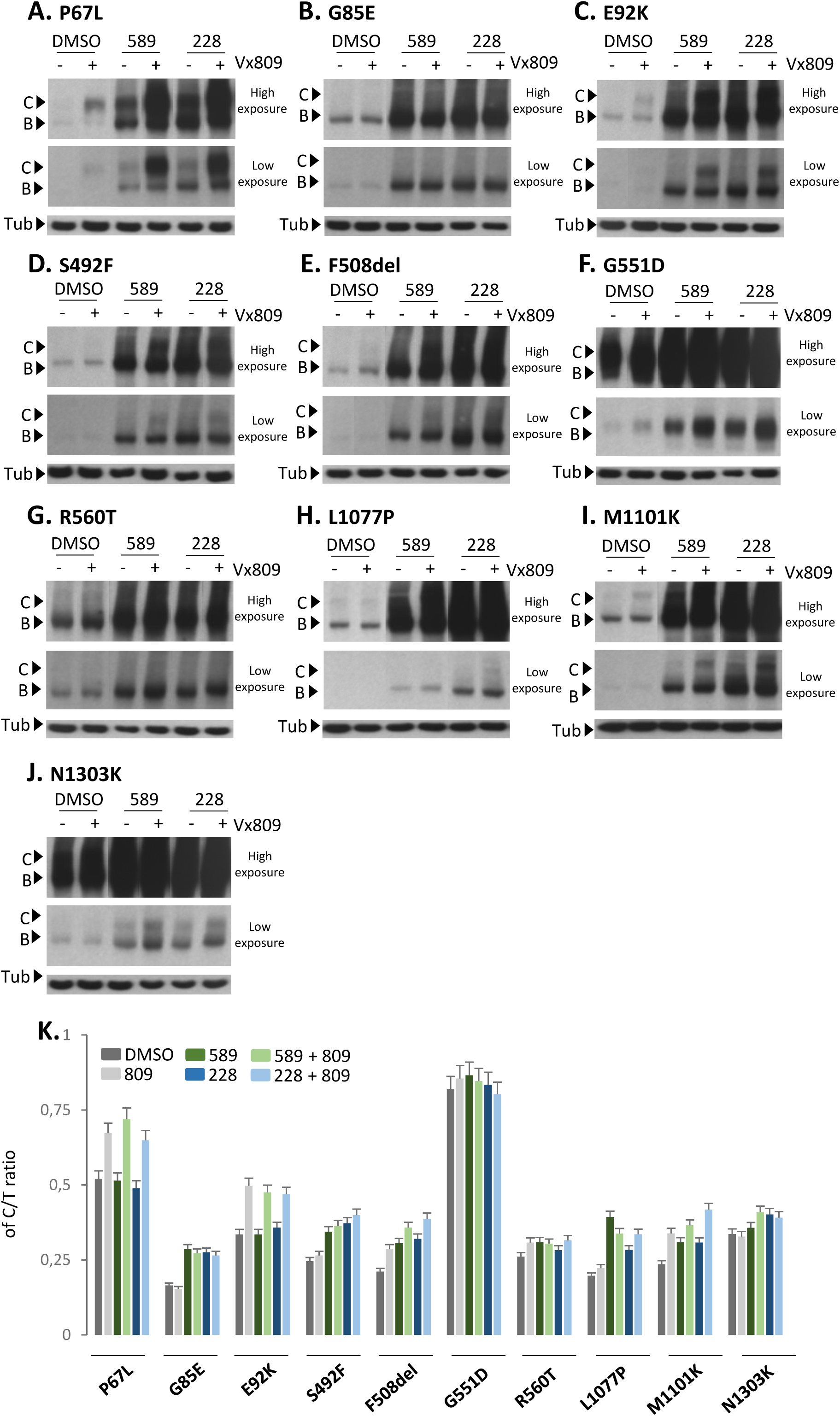
Effect of LBH-589 or FK-228 alone or in combination with Vx809 on CFTR2 variant trafficking. (**A to J**) Immunoblot analysis of CFTR expression following treatment of CFBE41o- transduced cells with 5 nM LBH-589 or 2.5 nM FK-228 with or without 3 µM Vx809 for P67L, G85E, E92K, S492F, F508del, G551D, R560T, L1077P, M1101K and N1303K variants respectively. (**K**) Quantification of the C/T ratio for each of the ten variants expressed as a % (mean ± SEM, n ≥ 3). Abbreviations: 589 (LBH-589) and 228 (FK-228).

While LBH-589 and FK-228 did not improve the C/T ratio for all variants tested, they did exhibit the ability to increase the amount of band C for all variants. To address if the corrected fraction of band C represented a functional pool of CFTR, we used the YFP quenching assay in transduced CFBE41o- null cells stably expressing the YFP-H148Q/I152L variant. LBH-589 and FK-228 were able to restore activity to a level exceeding 30% of WT CFTR YFP quenching for all CFTR2 variants tested (**Figure 8**). When we combined the HDACi treatment with Vx809, we further improved the activity of all variants. There were 2 striking observations made from these data. First, the combined treatment of HDACi + Vx809 for the P67L and E92K resulted in functional correction of these CF-causing variants to levels that exceed the activity seen with WT CFTR, suggesting that the HDACi-responsive PN modulations appear to be particularly permissive for these variants. Secondly, the HDACi treatment of the G551D variant corrected the functional defect associated with this class III variant to levels approaching that seen with WT CFTR.

**Figure 8.**
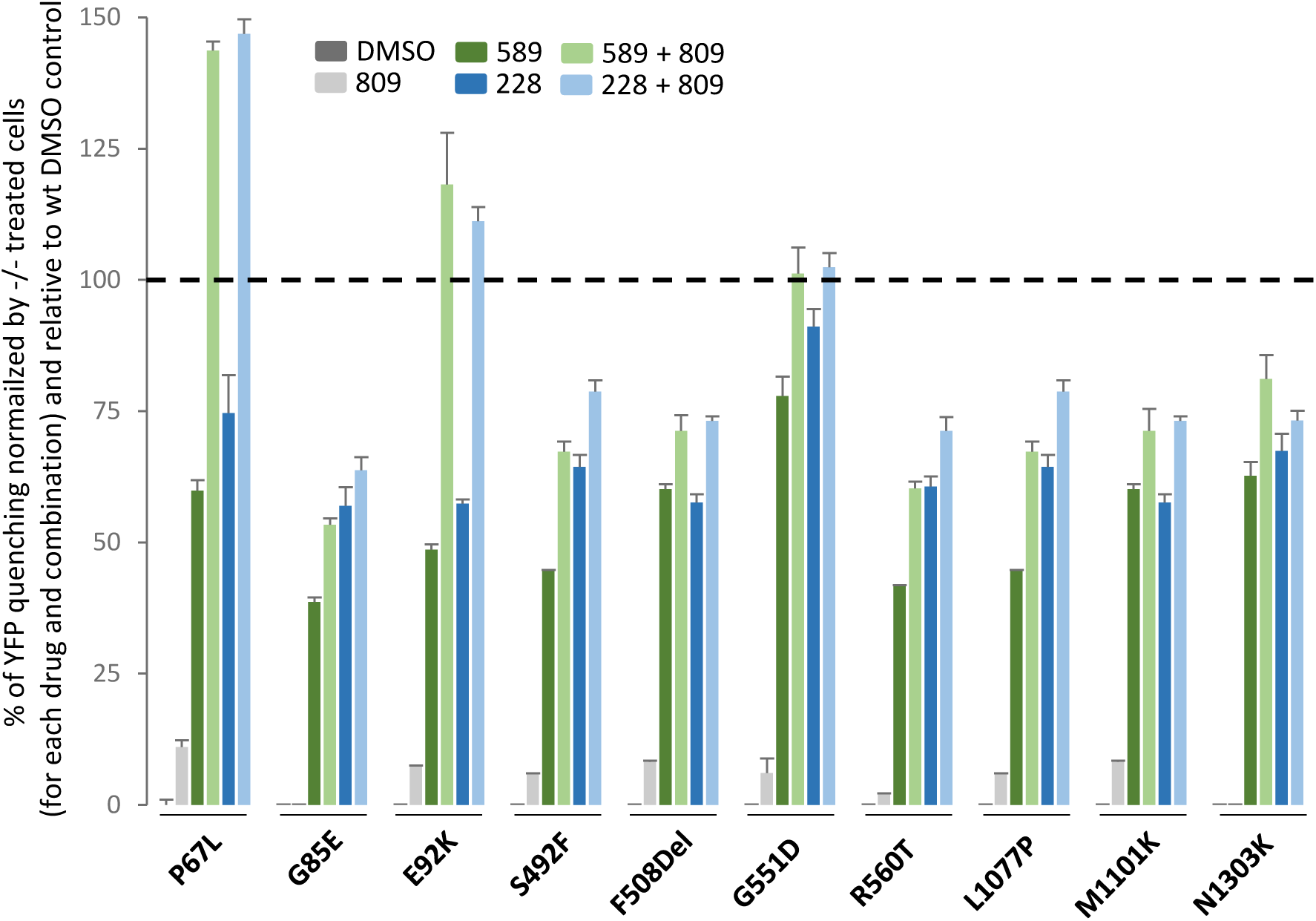
Effect of LBH-589 or FK-228 alone or in combination with Vx809 on CFTR2 variant activity. Quantification of YFP-quenching following treatment with 5 nM LBH-589 or 2,5 nM FK-228 with or without 3 uM Vx809 of CFBE41o- -YFP transduced cells with P67L, G85E, E92K, S492F, F508del, G551D, R560T, L1077P, M1101K and N1303K variants. Data are presented as a % of YFP quenching normalized by CFBE41o-YFP identically treated cells and relative to CFBE41o-YFP cells transduced with WT CFTR and treated with DMSO.

Taken as a whole, these data indicate that the HDACi-mediated correction of the F508del variant can be extend to other ER-restricted class II and III suggesting that the HDACi-mediated PN changes include alterations in the CFTR interaction network that impact the protein both in its early biogenesis as well as at PM where membrane localization, internalization and channel gating and function are also impacted.

## Discussion

There are currently a number of reports highlighting the beneficial impact of HDACi on abrogating the expression and aggregation of disease-causing protein variants (reviewed in [86, 107]) including the impact of Vorinostat on correcting the trafficking and functional defects associated with F508del-CFTR [90]. In the present study, we provide new evidence showing that LBH-589 and FK-228 can correct the trafficking defect associated with the F508del variant of CFTR leading to restoration of a functional chloride channel at the cell surface of both CFBE41o- and primary hBE cells. The HDACi-mediated correction of F508del-CFTR can synergize with the small molecule corrector, Vx809, to provide functional correction of this common CF-causing variant to a level approaching 40% of WT CFTR, a value that is predicted to provide significant therapeutic benefit in the clinic [102, 103]. Interestingly, while the HDACi, PXD-101, also showed corrector capability in the CFBE41o-model, it completely destabilized F508del CFTR in hBE cells, an effect which could not be overcome by Vx809 suggesting that this HDACi influences a different collection of HDAC activities than those seen in response to LBH-589 and FK-228 in hBE cells. Strikingly, the corrective properties of LBH-589 and FK-228 extend beyond the ability to correct F508del CFTR, as we show that they can also provide functional correction of other class II and III variants, both alone and in combination with Vx809, suggesting a more generalized mechanism of correction of CFTR variants. While we noted a generalized correction of the trafficking defect associated with the variants, we observe a differential effect on their functional correction. From the small cohort of variants tested, we identified 3 clusters of responses to HDACi: cluster A- class II variants exhibiting a modest functional correction; cluster B- class II variants exceeding WT activity; and cluster C- correction of the representative class III variant (**Figure 9**).

**Figure 9:**
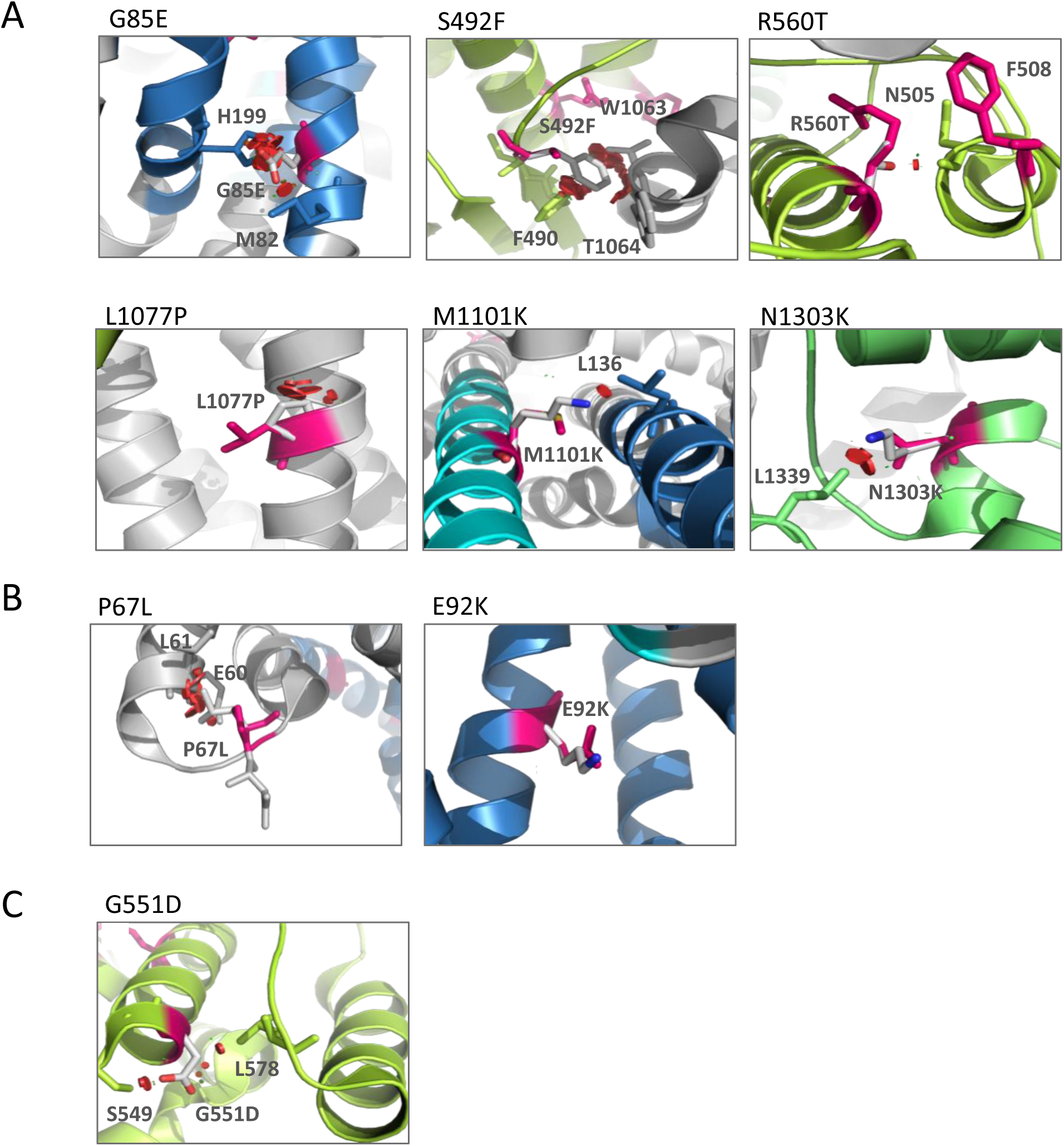
Structural modeling of CFTR2 variants. (**A to C)** Cryo-EM model of CFTR2 variants using PyMOL software (see Methods). Red shows impact of the mutant residue on other residues. Rotomers are predicted to minimize impact against other residues. (**A**) Cluster A: G85E, S492F, G551D, R560T, L1077P, M1101K and N1303K. (**B**) Cluster B: P67L and E92K variants. (**C**) Cluster C: G551D variant.

Cluster A variants include G85E, S492F, F508Del, R560T, L1077P, M1101K and N1303K (**Figure 9A**), which exhibit a modest correction in response to HDACi. The G85E variant, which is localized in TMD1 (**Figure 9A**), represents the least responsive variant tested. The substitution of glycine for glutamic acid at this position drastically impacts CFTR structure both locally, given the interaction of G85 with M82, and globally, given the interaction of G85 with H199. These data are in agreement with the observations that G85E is a severe class II variant disrupting the ER targeting, integration and topology of the TMD1 [108, 109], which cannot be overcome with low temperature nor with well-established correctors such as C4, C18 and Vx809 [110]. While the impact of Vx809 alone was hard to detect, we observed a synergistic correction of the variants in response to HDACi + Vx809, an effect that is most pronounced with LBH-589. These data suggest that HDACis, are able to make adjustments to the cellular environment, influenced by epigenomic changes [90], which allow a global correction of previously misfolded class II CFTR variants. The severity of the defect seen with specific variants can generate an impediment to correction, suggesting that they may lie outside the tolerance window for functional structure correction [2].

The most responsive variant is the cluster B, P67L variant, which is localized in the N-terminal region of the CFTR polypeptide near the TMD1 (**Figure 9B**). An *in silico* analysis of the impact of substituting the proline at position 67 with a leucine (**Figure 9B**) suggests that the mutation would impact the structure of CFTR due to a predicted clash of L67 with both E60 and L61 thereby limiting flexibility that may be essential for generating the functional fold. The P67L variant was initially classified as a class IV gating variant [111], suggesting that it exhibited a significant amount of trafficking at steady state. However, recent evidence has shown that the P67L mutation is in fact a class II variant exhibiting weak trafficking, yet is highly responsive to Vx809 [112]. Our data, presented herein are in agreement with these observations, revealing that P67L CFTR is a class II variant which exhibits functional correction in response to Vx809 that synergizes with LBH-589 and FK-228 to restore activity to a level seen with the WT. Considering that the P67L variant is characterized by an elevated C/T ratio, we would predict that LBH-589 and FK-228 (in combination Vx809) could provide significant benefit to 10 other variants which achieve a C/T ratio of at least 30% including R74W,R75Q, I336K, S341P, R347P, S549R, D579G, D614G, S945L and R1070W [4].

The second most responsive variant to HDACi is the cluster B, E92K variant, which localizes to the first membrane helix of the of the TMD1 (**Figure 9B**). Analysis of the impact of substituting the glutamic acid at position 92 with a lysine (**Figure 9B**) suggests that the mutation would not severely impact the structure of CFTR- accounting for its high responsiveness to HDACi and Vx809. Overall, the HDACi-mediated response of cluster B variants (**Figure 9B**) compared to cluster A (**Figure 9A**) suggests that cluster B mutations present a less severe disruption of the CFTR 3D structure making them more sensitive to epigenetic changes that are incurred in response to alterations in the cellular acetylation status mediated through HDACis. We also observed that HDACi treatment was sufficient to restore activity to the class C variant G551D CFTR (**Figure 9C**) found at the cell surface to WT levels, suggesting that the epigenetic changes generated by LBH-589 and FK-228 are able to restore key interactions of the G551D polypeptide with the gating machinery controlling channel conductance.

In general, we now appreciate that HDACi alter the acetylation equilibrium of both histones (regulating transcriptional events) and non-histone proteins, an effect highlighted by the dynamic acetylome, which included proteostasis components [1, 25–31] regulating the central heat shock response (HSR) [32] and unfolded protein response (UPR) [33, 34], programs responsible for recovery from protein folding stress such as the maladaptive stress caused by CFTR misfolded variants [24, 35–41]. By globally adjusting the cellular environment, HDACi could be adjusting interacting pathways that abrogate maladaptive states that arise in response the expression of disease-associated variants thereby creating a more permissive folding environment. While most efforts using HDACi have focused on killing cancer cells, a number of approaches have now demonstrated their value in human disease. For example, LBH-589 has been reported to rescue sickle cell disease (SCD) [110] and its clinical benefits are under investigation. While the toxicity of these compounds has been well documented, they have been shown to be sufficiently well tolerated by patients even at doses toxic to cancer cells, which far exceed the doses at which we see functional correction of CF-causing variants, suggesting that a therapeutic index may exist for FDA-approved HDACi to provide significant benefit to a broad spectrum CF patients.

## Materials and Methods

### Cell culture and treatment

Null (-/-) CFBE41o-, CFBE41o- expressing F508del CFTR (F508del CFBE41o-) or Wild-Type (WT) CFTR (WT-HBE41o-) were cultured as previously described [90]. CFBE41o-, F508del CFBE41o- and WT-HBE41o- were stably transfected with pcDNA3.1 containing YFP-H148Q (kindly provided by Dr. L. Galietta, Telethon University for Genetics and Medicine, Puzzuoli, IT) using Lipofectamine 3000 (Invitrogen) according to the manufacturer’s instructions. CFBE41o-YFP, F508del CFBE41o-YFP and WT HBE41o-YFP were selected with 0.75 mg/ml G418 and then sorted using FACS. The stable cell lines could be passed at least 15 times without a decrease in YFP-H148Q fluorescence. CFBE41o-YFP, F508del CFBE41o-YFP and WT HBE41o-YFP were cultured in the same media as the YFP non-expressing cells with addition 0.75 mg/ml G418. Cells were treated at the indicated concentration of belinostat (Toronto Research Chemicals, Inc), panobinostat (LC laboratories) and romidepsin (Toronto Research Chemicals, Inc) in complete growth media and incubated at 37°C, 5% CO2 for 24 h. 3 µM of Vx809 was used where indicated.

### LDH cytotoxicity assay

Following the treatment of F508del CFBE41o- with 10 pM to 10 µM PXD-101, LBH-589 or FK-228, cytotoxicity of the three compounds were monitored following the manufacturer instructions of the LDH cytotoxicity assay kit (Pierce).

### Adenoviral transduction

CFTR2 variants coding sequence were PCR amplified using forward (TCATGGTACCATGCAGAGGTCGCCT) and reverse (GCTGCTCGAGCTAAGCGTAATCTGGAACATCGTATGGGTAAAGCCTTGTATCTTG) primers introducing a HA-TAG at the end of the sequence and pBl-CMV2-CFTR plasmids DNA (kindly provided by P. Thomas, University of Texas Southwestern) as template. The PCR fragments were digested with KpnI and XhoI and cloned into pENTR1A shuttle vector. pENTR1A-CFTR plasmids were sequenced and recombined into pAD-CMV-V5-DEST using LR-clonase II (Invitrogen). pAD-CMV-CFTR plasmid were then sent to ViroQuest for adenovirus production [42, 113]. 80% confluent CFBE41o- were transduced for 5 h in OPTI-MEM in the presence of 10 µg/ml of polybrene (EMD Milipore Corp) with adenovirus carrying CFTR at a multiplicity of infection of 200. Then cells were washed with PBS and fed with regular medium for 24 h prior to treatment. Kinetics of CFTR mRNA and protein level for wild-type and F508del were performed to optimize the efficiency of CFTR expression. To ensure a similar mRNA level of the different CFTR2 variant after adenovirus transduction, qRT-PCR was performed with no significant differences detected.

### qRT-PCR

qRT-PCR was performed using the iScript One-Step RT-PCR kit with SYBR green (Bio-Rad). RNA was standardized by quantification of beta-glucuronidase (GUS) mRNA, and all values were expressed relative to GUS. Forward (GTGGCTGCTTCTTTGGTTGT) and reverse (CGAACTGCTGCTGGTGATAA) primers were used as indicated.

### Immunoblotting

CFBE41o- cells lysates were prepared in 50 mM Tris-HCl, 150 mM NaCl, 1% Triton X-100 and 20 µg of total protein were separated on an 7% SDS-PAGE. Proteins were transferred to nitrocellulose and probed with 3G11 antibody for CFTR detection and rat secondary antibody, and detected by chemiluminescence. The same procedure was followed for primary F508del/F508del hBE cells lysate with modifications. Benzonate was added to the lysate buffer at a final concentration of 0.025 units/µl and 40 µg instead of 20 µg of total protein were used for SDS-PAGE.

### YFP quenching assay

CFBE410- cells expressing F508del CFBE41o-YFPwere treated with indicated drug. Following treatment, cells were stimulated with a final concentration of 10 µM Forskolin and 50 µM Genistein for 15 min prior to addition of PBS + NaI (replacement of NaCl with 137 mM NaI). Fluorescence was monitored every second for a total of 30 sec (3 sec prior to addition of NaI and 27 seconds after addition of NaI) using a Synergy H1 hybrid reader (Biotek). As a negative control, identical experiments were performed with CFBE41o- YFP cells not expressing F508del CFTR.

### Ussing chamber conductance assay cell culture

Primary F508del/F508del hBE cells were cultured on permeable supports as described previously [101]. Short-circuit currents (Isc) were measured as previously described [101].

## Acknowledgements

We thank P. Thomas (UT Southwestern Medical School) for providing vectors encoding CFTR2 mutants and Dr. L. Galietta (Telethon Institute of Genetics and Medicine, IT) for providing the vector encoding YFP-H148Q. Support provided by the National Institutes of Health Grants HL095524 and DK051870 to WEB, and by a Cystic Fibrosis Foundation Therapeutics (CFFT) Fellowship to Frédéric Anglès.

## Supplementary Figure Legends

**Figure S1.**
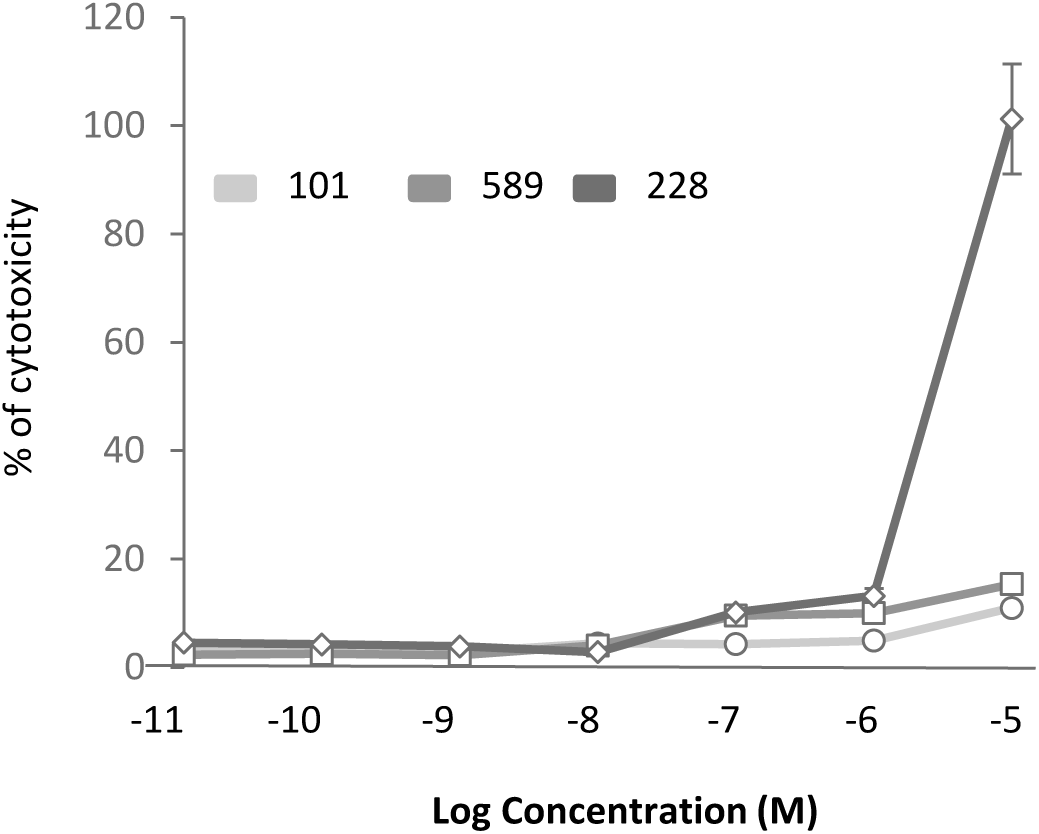
Cytotoxicity of PXD-101, LBH-589 and FK-228 on CFBE41o-YFP and F508del CFBE41o-. Quantification of PXD-101, LBH-589 and FK-228 cytotoxicity following treatment of F508del CFBE41o- cells using the LDH cytotoxicity assay kit (mean ± SEM, n ≥ 3).

**Figure S2.**
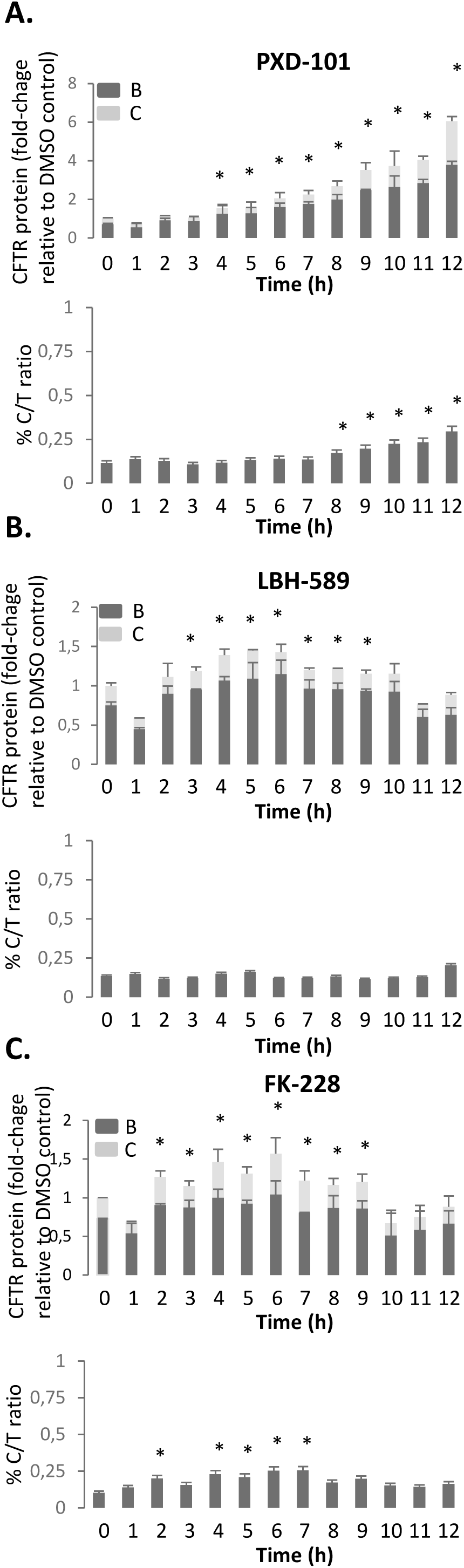
Quantification of F508del CFTR rescue following PXD-101, LBH-589 or FK- 228 treatment during the first 24 h. (**A, B & C**) Quantification analysis of CFTR expression following treatment of F508del CFBE41o- cells with 2 µM PXD-101, 10 nM LBH-589 and 5 nM FK-228 respectively. Data are presented in upper panel as fold-change relative to DMSO treatment (mean ± SEM, n ≥ 3). Lower panel present quantification of C/T ratio expressed as a % (mean ± SEM, n ≥ 3). In all panels, * indicates significant difference (p < 0.05) relative to DMSO treatment as determined by two-tailed t-test.

**Figure S3.**
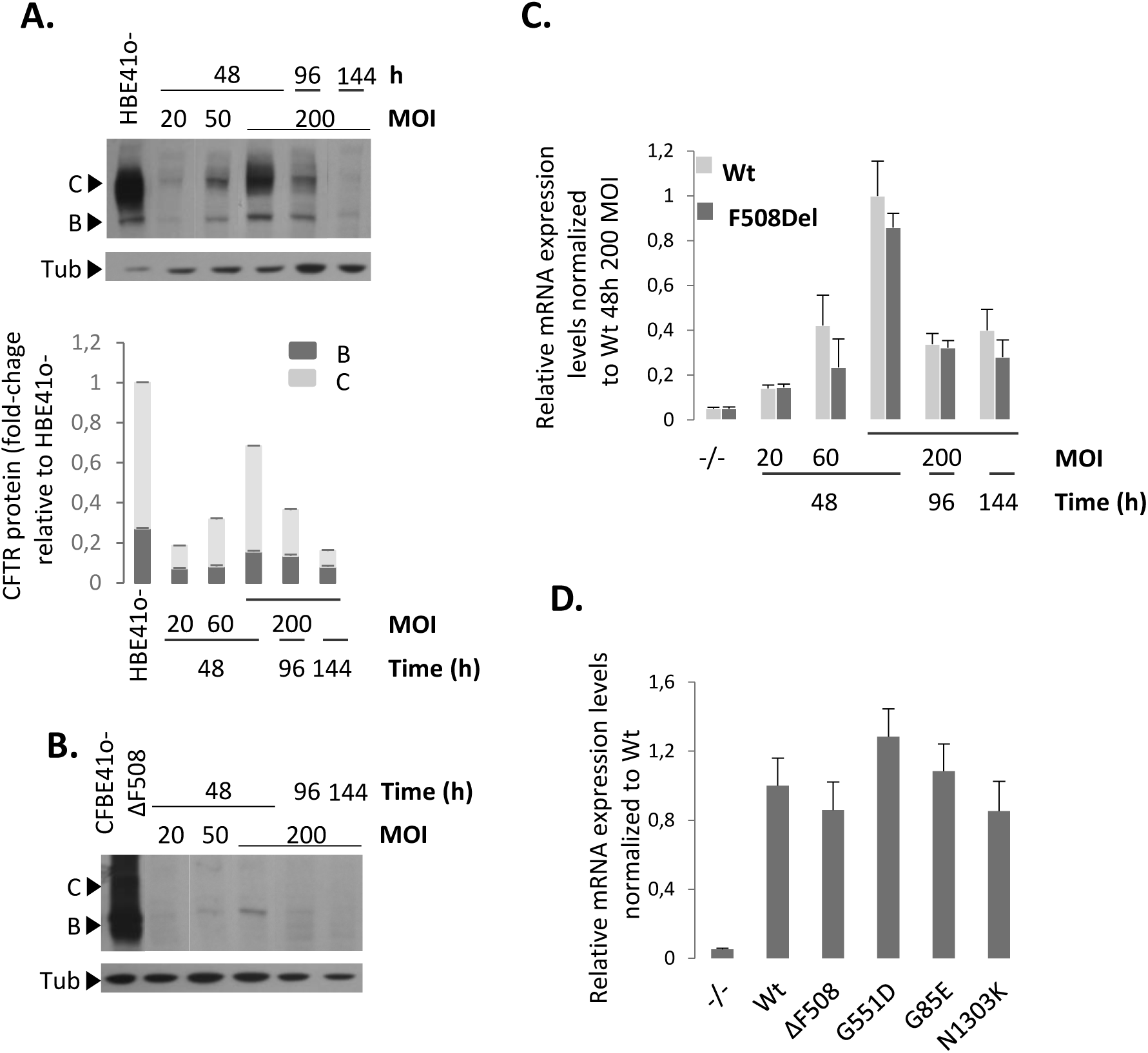
Optimization and validation of CFBE41o- CFTR-adenoviral transduced cell model. (**A**) Immunoblot analysis (upper) and quantification (lower) of WT CFTR expression in CFBE41o- cells following transduction with 20, 50 or 200 multiplicity of infection (MOI) after 48, 96 and 144 hours. Data are presented as fold-change relative to DMSO treatment (mean ± SEM, n ≥ 3). (**B**) Immunoblot analysis of F508del CFTR expression in CFBE41o- cells following transduction with 20, 50 or 200 multiplicity of infection (MOI) after 48, 96 and 144 hours. (**C**) Quantification of CFTR F508del or WT mRNA level in CFBE41o- cell following transduction with 20, 50 or 200 MOI after 48, 96 and 144 h. mRNA was standardized by quantification of beta-glucuronidase (GUS) mRNA, and all values were expressed normalized to GUS (mean ± SEM, n ≥ 3). (**D**) Quantification of CFTR WT, F508del, G551D, G85E and N1303K mRNA level following transduction with 200 MOI after 48, 96 and 144 hours. mRNA was standardized by quantification of GUS mRNA, and all values were expressed normalized to GUS and relative to WT mRNA level (mean ± SEM, n ≥ 3).

**Figure S4.**
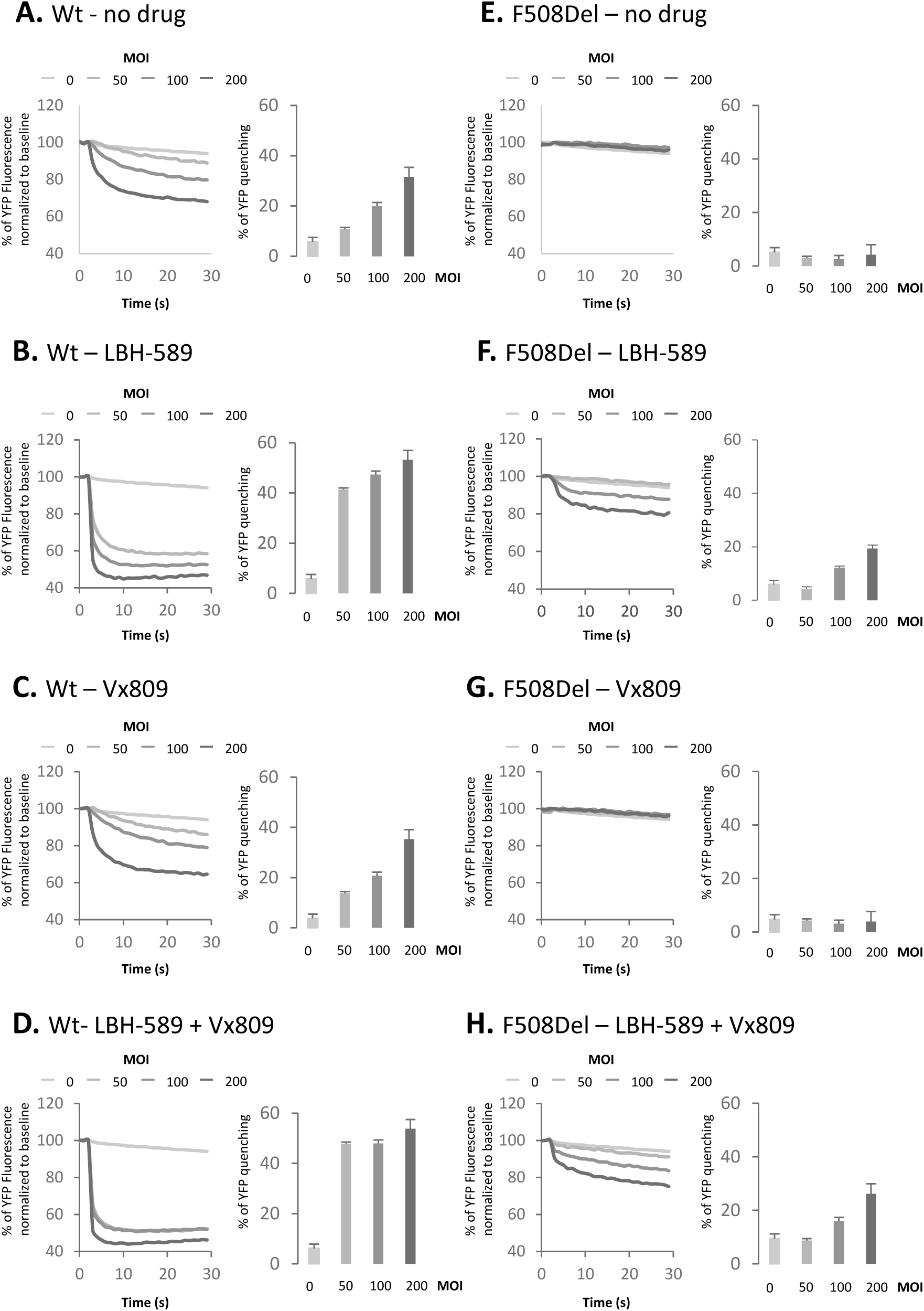
Optimization and validation of CFBE41o-YFP CFTR-adenoviral transduced cell model. Representative FLIPR traces (left) and quantification (right) of YFP-quenching following transduction of CFBE41o-YFP cells with WT CFTR (**A to D**) or F508del CFTR (**E to H**), with 50, 100 and 200 MOI and treated with DMSO (**A & E**), 5 nM LBH-589 (**B & F**), 3 µM Vx809 (**C & G**) or a combination of 5 nM LBH-589 and 3 µM Vx809 (**D & H**). Data are presented as % normalized to baseline (mean ± SEM, n ≥ 3).

